# DARTsort: A modular drift tracking spike sorter for high-density multi-electrode probes

**DOI:** 10.1101/2023.08.11.553023

**Authors:** Julien Boussard, Charlie Windolf, Cole Hurwitz, Hyun Dong Lee, Han Yu, Olivier Winter, Liam Paninski

## Abstract

With the advent of high-density, multi-electrode probes, there has been a renewed interest in developing robust and scalable algorithms for spike sorting. Current spike sorting approaches, however, struggle to deal with noisy recordings and probe motion (drift). Here we introduce a modular and interpretable spike sorting pipeline, **DART**sort (**D**rift **A**ware **R**egistration and **T**racking), that builds upon recent advances in denoising, spike localization, and drift estimation. DARTsort integrates a precise estimate of probe drift over time into a model of the spiking signal. This allows our method to be robust to drift across a variety of probe geometries. We show that our spike sorting algorithm outperforms a current state-of-the-art spike sorting algorithm, Kilosort 2.5, on simulated datasets with different drift types and noise levels. Open-source code can be found at https://github.com/cwindolf/dartsort.

## 1 Introduction

Recently, there has been a surge in the development of high-density micro-electrode arrays (MEAs) which are able to record from large populations of neurons with high temporal resolution (*Frey et al., 2010; Ballini et al., 2014; Lopez et al., 2016; Jun et al., 2017; Angotzi et al., 2019; Juavinett et al., 2019; Steinmetz et al., 2021; Trautmann et al., 2023*). Many of these devices can record activity from multiple brain areas concurrently, promising to improve our understanding of distributed neural computation (*Chung et al., 2019*).

Spike sorting is a critical first step in the analysis of single-neuron activity. A key assumption in spike sorting is that neurons have a characteristic spatiotemporal footprint (i.e. template) that is dependent on the morphology, position, and orientation of the cell (*Gold et al., 2006*). Using this assumption, spike sorting algorithms can demix and assign spikes to their putative neurons (*Lewicki, 1998*). Numerous spike sorting algorithms have recently been proposed to take advantage of multi-electrode recordings (*Pillow et al., 2013; Garcia and Pouzat, 2015; Rossant et al., 2016; Pachitariu et al., 2016; Chung et al., 2017; Hilgen et al., 2017; Yger et al., 2018; Diggelmann et al., 2018; Chaure et al., 2018; Lee et al., 2020; Steinmetz et al., 2021; Chen et al., 2021*).

A major unsolved challenge for spike sorting in high-density MEAs is probe motion (drift) relative to the brain, which induces nonstationarities in spike shapes and amplitudes (*Buccino et al., 2022*). These nonstationarities break the assumption that neurons have a characteristic extracellular signature and can lead to numerous spike sorting issues, including units disappearing (or appearing) over time, or neurons being split into multiple units (oversplitting) (*IBL, 2022*).

Two current approaches to correcting for drift have been pursued: (1) empirically updating each unit’s template after a fixed amount of time (*Yger et al., 2018; Lee et al., 2020*) and (2) estimating the motion of the probe during the recording and then performing a spatial resampling/interpolation of the raw data according to this estimate (*Steinmetz et al., 2021*). The first method works best if drift is gradual (so that changes in the template are slow and can be tracked easily), but does not take advantage of the fact that drift is highly correlated across multiple nearby cells. The second method, while widely used, is suboptimal for several reasons. First, it assumes that the template of each cell is fixed following the resampling/interpolation step; this assumption typically does not hold due to both imperfections in registration/interpolation and also due to slow changes in the interface between individual neurons and the MEA over time. Second, spatial resampling is not able to correctly infer signal near the edge of the probe since this can require an extrapolation beyond the probe boundary. Finally, as interpolation/resampling relies on a spatial averaging of the signal, it generally reduces the signal-to-noise ratio (SNR).

A recent alternative arises from new developments in spike localization for high-density MEAs (*Jun et al., 2017; Hilgen et al., 2017; Hurwitz et al., 2019; Boussard et al., 2021*). After estimating the source location of the detected spikes, it is possible to construct a precise map of the drift over time using a motion estimation algorithm which is able to account for fast time scale drifts (*Varol et al., 2021; Windolf et al., 2022*). We postulate that a spike sorter which incorporates this precise drift map into its model of spiking activity could achieve the “best of both worlds,” combining the strengths of the current two approaches described above and therefore leading to better performance overall.

In this paper, we present a modular spike sorting pipeline, DARTsort, that incorporates state-of-the-art spike localization (*Boussard et al., 2021*) and drift estimation (*Windolf et al., 2022; Garcia et al., 2023*) to explicitly model the neurons’ spatiotemporal signature as a function of location and time. Rather than modeling each unit with a single template, a set of spatially “superresolved” templates is constructed by binning each unit into sub-units across the depth of the probe in order to model the effect of drift. Then, these superresolved templates (and the corresponding drift map) are directly incorporated into the template matching step. (In other words, we register the templates to the drifting data, rather than attempting to register the raw data as in *Steinmetz et al*. (*2021*).) This allows DARTsort to resolve fast time scale drift which will cause sudden changes in each unit’s template. Along with our novel drift correction approach, we also present a number of improvements to denoising, spike detection, spike featurization and clustering, and unit splitting and merging, all of which further boost the performance of our algorithm. We show that DARTsort outperforms Kilosort 2.5 on simulated data across multiple drift regimes and noise levels. As our codebase is highly modular, many of our processing steps have already been incorporated into SpikeInterface (*Buccino et al., 2020*), an open-source Python frame for spike sorting. Overall, we demonstrate that DARTsort is a scalable and robust spike sorting solution for high-density MEAs. The code for DARTsort can be found at https://github.com/cwindolf/dartsort.

## 2 Overview of sorting algorithm steps

At a high level, the algorithm proceeds as follows (shown more formally in Algorithm 1; see also Figure 1). Similarly to clustering algorithms which alternate between updating cluster parameters and assigning data points to individual clusters, the algorithm will alternate between updating a set of candidate templates for each unit and updating which spikes are assigned to each unit.

### Algorithm 1

Main sorting routine.

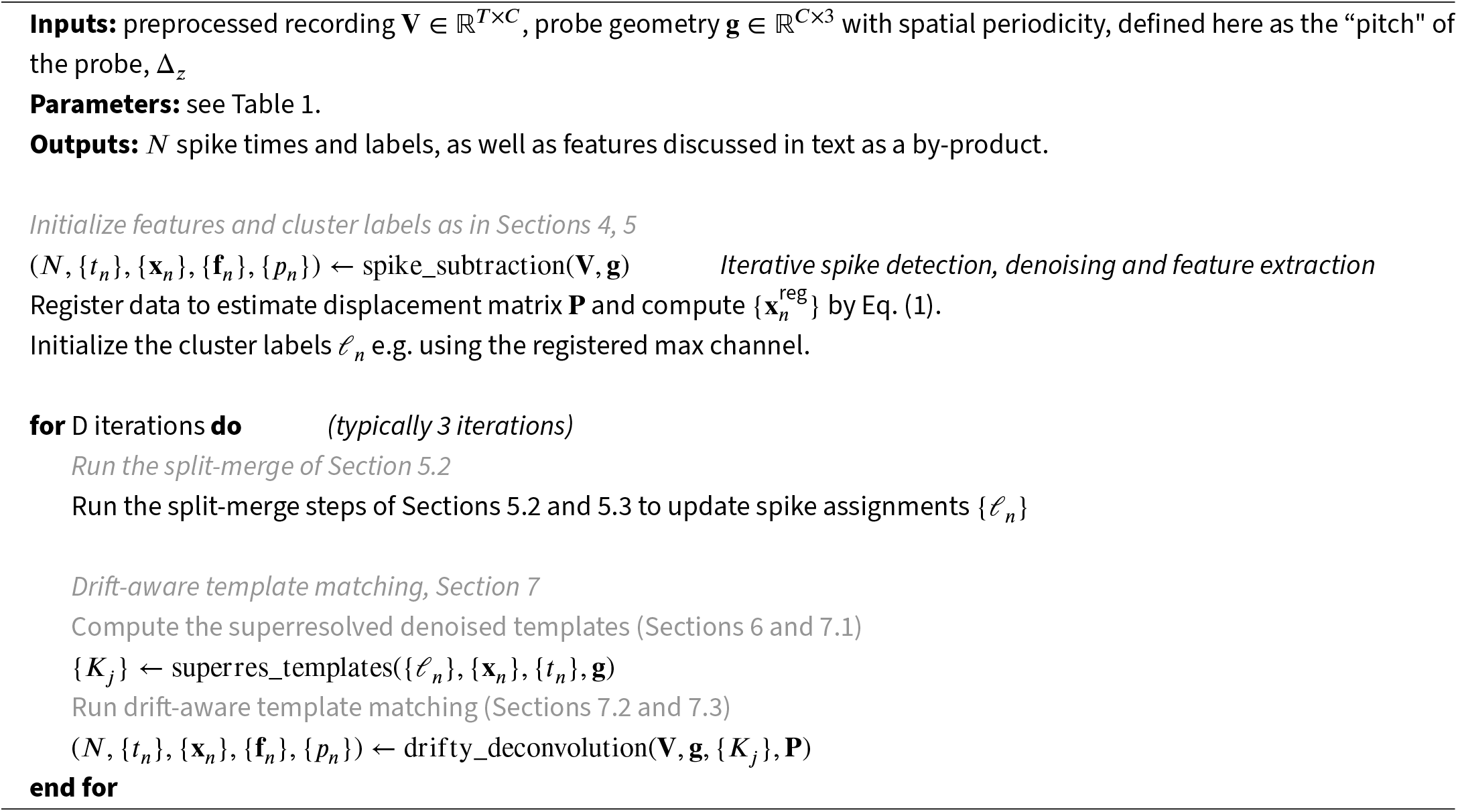

**Figure 1.**
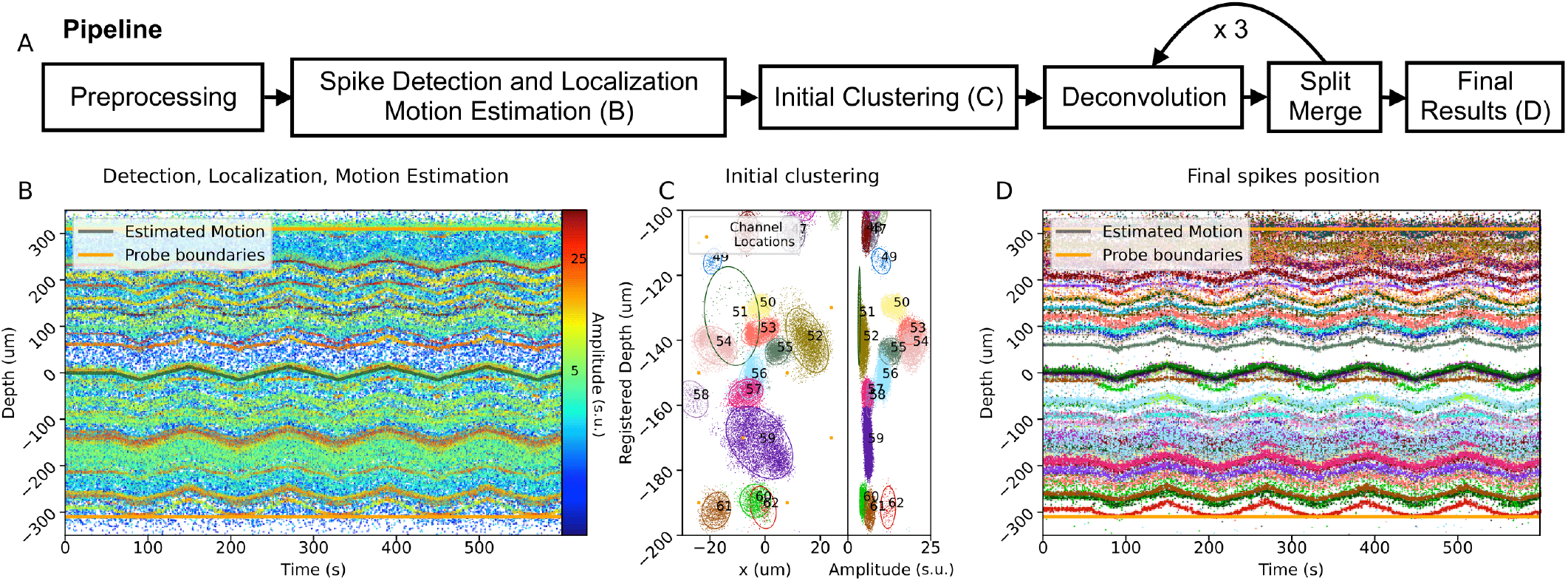
Overview of the DARTsort pipeline and results on simulated data with rigid drift. **(A)** Overview of each processing step in DARTsort. **(B)** After preprocessing the recording, spikes are detected, denoised and localized, before the motion is estimated for the recording **(C)**. An initial clustering is run using extracted and registered spike features. **(D)** Alternating steps of the drift-aware deconvolution and split-merge steps are run to find stable units even in the presence of drift.

To initialize the alternating algorithm, a first detection step (Section 4) is run, yielding *N* spikes with detection times *t*_*n*_. For each spike, the peak-to-peak (ptp) amplitude *p*_*n*_ and source location **x**_*n*_ = (*x*_*n*_, *y*_*n*_, *z*_*n*_) are computed, where *z*_*n*_ represents the depth of the spike along the probe, *x*_*n*_ its lateral position, and *y*_*n*_ its position orthogonal to the plane of the probe. All these features are input into a motion estimation algorithm (DREDge; *Windolf et al*. (*2022*)) to estimate the depth motion traces P ∈ ℝ^*T* ×*D*^ in *D* depth bins across the probe. This motion estimate is used to shift each spike to its ‘registered’ depth position. Finally, cluster assignments are initialized using the clustering method described in Section 5.1. Initial templates are computed by denoising the median of the waveforms in each cluster (Section 6).

Next, a split-merge and template matching step are alternated. First, a split-merge step is applied to the current cluster assignments *ℓ*_*n*_ (Section 5.2). In the split step, temporal PCA features **f**_*n*_ are extracted for each cluster on a subset of channels around the cluster’s max channel and are transformed to account for drift (Section 4.3). Two PCA features are then combined with the localization features and clustered with HDBSCAN (*McInnes et al., 2017*). This split step is run recursively (*Swindale and Spacek, 2014*), re-fitting the per-cluster PCA embeddings in each iteration and applying HDBSCAN until no new splits are found. Next, a distance is computed between all pairs of shifted, scaled, and normalized templates and a merge is performed when this distance is small.

After the split-merge step finishes updating the cluster assignments *ℓ*_*n*_, the spikes in each unit are binned in a drift-aware manner (Section 7.1), leading to a set of spatially superresolved cluster assignments. A denoised template (Section 6) is computed for each superresolved group. A template matching procedure (Section 7.3) is then carried out to update each template’s set of spikes, i.e., the spike count *N*, the spike times *t*_*n*_ and labels *ℓ*_*n*_, the localization features **x**_*n*_, and the temporal PCA subspace and loadings **f**_*n*_. This updated spiking point process can be fed back into the split merge and these two steps can be alternated until convergence or until a computational budget is reached.

**Table 1.**
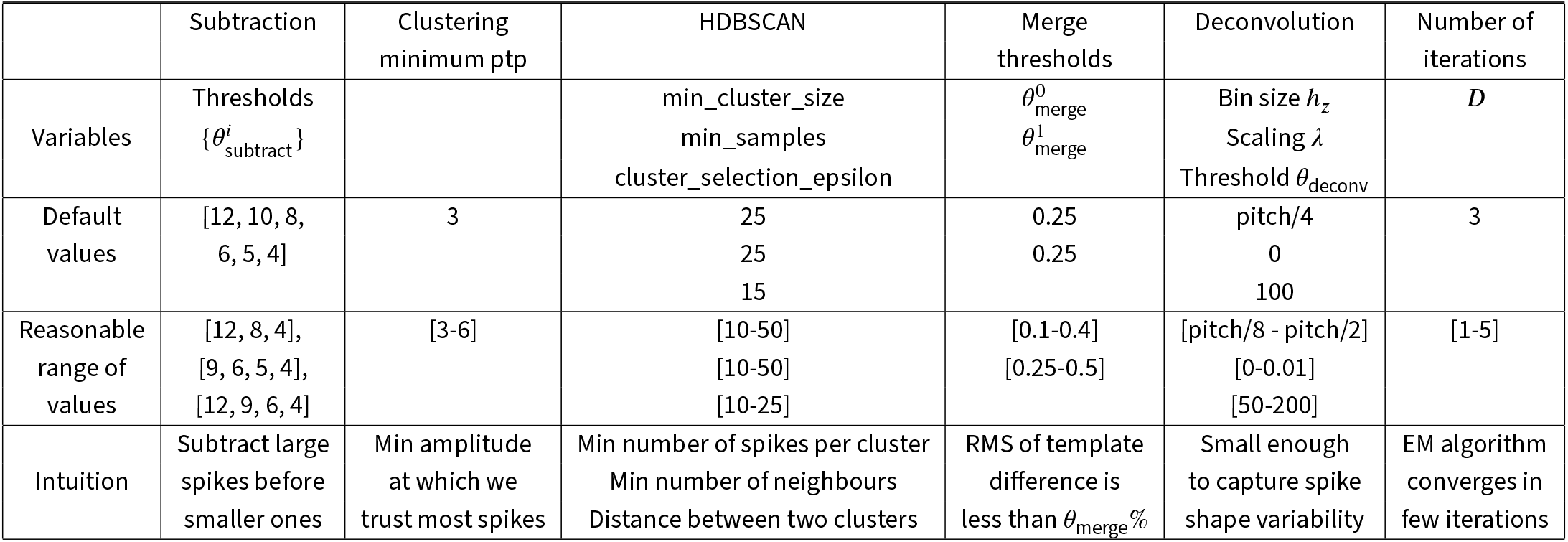
List of all parameters used in DARTsort. For each parameter, the default value is provided (second row) as well as a reasonable range of values (third row).

|  | Subtraction | Clustering<br>minimum ptp | HDBSCAN | Merge<br>thresholds | Deconvolution | Number of<br>iterations |
| --- | --- | --- | --- | --- | --- | --- |
| Variables | Thresholds<br>$\{\theta_{\text{subtract}}^i\}$ | | min_cluster_size<br>min_samples<br>cluster_selection_epsilon | $\theta_{\text{merge}}^0$<br>$\theta_{\text{merge}}^1$ | Bin size $h_z$<br>Scaling $\lambda$<br>Threshold $\theta_{\text{deconv}}$ | $D$ |
| Default<br>values | [12, 10, 8,<br>6, 5, 4] | 3 | 25<br>25<br>15 | 0.25<br>0.25 | pitch/4<br>0<br>100 | 3 |
| Reasonable<br>range of<br>values | [12, 8, 4],<br>[9, 6, 5, 4],<br>[12, 9, 6, 4] | [3-6] | [10-50]<br>[10-50]<br>[10-25] | [0.1-0.4]<br>[0.25-0.5] | [pitch/8 - pitch/2]<br>[0-0.01]<br>[50-200] | [1-5] |
| Intuition | Subtract large<br>spikes before<br>smaller ones | Min amplitude<br>at which we<br>trust most spikes | Min number of spikes per cluster<br>Min number of neighbours<br>Distance between two clusters | RMS of template<br>difference is<br>less than $\theta_{\text{merge}}\%$ | Small enough<br>to capture spike<br>shape variability | EM algorithm<br>converges in<br>few iterations |

## 3 Preprocessing

For preprocessing, the pipeline recommended by IBL (*IBL, 2022*) is utilized. First, a high-pass filter is run on the data with a cutoff frequency of 300Hz. Second, the data is standardized by dividing each channel by the median absolute deviation of its noise. Third, an Analog to Digital converter (ADC) is run. Finally, a ‘destriping’ step is run on the data by using a spatial filter to remove artifacts caused by ADC lag.

## 4 Initial detection and featurization by neural-net aided blind deconvolution

Given a set of templates obtained from previous spike sorting results, a neural network is trained to denoise spikes. The network takes, as input, temporally-aligned, single-channel spike waveforms and attempts to output the corresponding templates. The spike waveforms are created by corrupting the set of templates with various naturalistic noise sources. The procedure described in (*Lee et al., 2020*) is followed for this step. Importantly, one of the data augmentations consists of colliding two templates, which allows the network to be robust to spikes that overlap with other spikes.

Spike denoising can be improved by subtracting away big spikes before detecting and denoising small spikes. After detection of candidate spiking events (with amplitude larger than some threshold 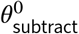, the neural network is used to denoise the spikes.

To improve the pipeline’s robustness to neural network errors, these spike waveforms are further denoised with PCA, fit and applied on each channel independently, and rescaled on each channel such that the peak-to-peak amplitude of each channel decreases as the distance to the peak channel grows. The denoised waveforms are then subtracted from the recording, leaving behind a residual; as in the template matching procedure below, spikes whose subtraction would fail to decrease the norm of the residual by an amount larger than a fixed threshold are rejected. This procedure is repeated greedily over a set of decreasing thresholds 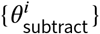 See Fig. 2 for an illustration.

**Figure 2.**
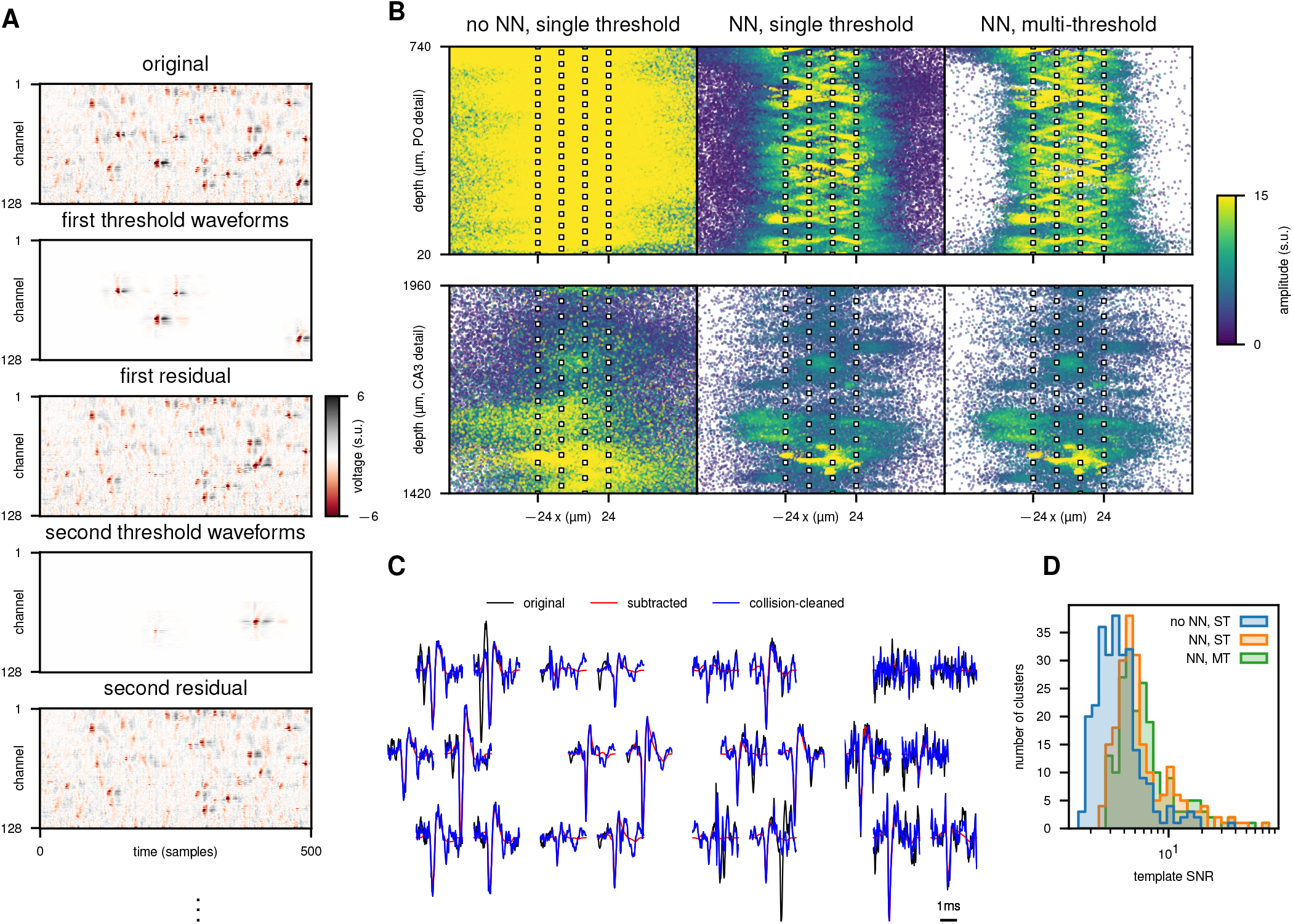
Overview of the initial detection and featurization step applied to a 5 minute segment of an IBL Neuropixels recording (*IBL et al., 2022*). **(A)** shows the iterative subtraction procedure on an illustrative subset of data. Starting from the original (preprocessed) recording, detections above a threshold are denoised and subtracted to yield a residual. This step is repeated with successively lower thresholds. **(B)** shows the utility of denoising and iterative subtraction. Spike localizations are shown in two brain regions (rows) with no denoising or subtraction, denoising without subtraction, and denoising with subtraction (left to right), colored by amplitude. Localizations of high-amplitude (yellow) spikes have large variance without neural net denoising; iterative subtraction resolves a cloud of noisy localizations for small-amplitude (purple) spikes. This procedure’s ability to resolve collisions on specific waveforms is shown in **(C)**, where original (preprocessed) waveforms, the waveforms subtracted during the iterative subtraction-and-denoise procedure, and the collision-cleaned waveforms (sum of subtracted and final residual) are overlaid. Panel **(D)** shows the cluster signal-to-noise ratio (SNR) obtained after clustering the resulting spike localizations following the method described in Section 5.1.2. The cluster SNR measures how close the template is to its cluster’s spikes, and is computed as the standard deviation of the dot product between a template and its corresponding spikes, divided by the standard deviation of the dot product between the template and random snippets of data. Templates were obtained after running our initial clustering on the spike localization features (Section 5). Using the neural network denoiser and subtraction increases SNR.

After this iterative detect-denoise-subtraction procedure finishes, a final residual remains. A set of collision-cleaned waveforms **w**_*n*_ are created by adding the final residual back to the subtracted and denoised waveforms. As other waveforms have been subtracted to obtain the final residuals, these waveforms are free from collisions. A final set of denoised waveforms can then be produced by passing the collision-cleaned waveforms **w**_*n*_ through the denoiser.

In the absence of a pretrained neural network, the user can choose to skip the subtraction routine. In this case, a typical peak detection pipeline is utilized where waveforms are extracted, centered on the detected peaks, and denoised. These waveforms are used in place of the collision-cleaned waveforms **w**_*n*_. For recordings with few collisions, the final waveforms will be similar to the collision-cleaned waveforms.

### 4.1 Spike feature extraction

A set of low-dimensional features is computed for each spike using the denoised waveforms. These features include the localizations **x**_*n*_, ptps **p**_*n*_, the spread *s*_*n*_ defined as 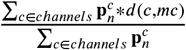 (with *d*(*c, mc*) denoting the Euclidean distance between channel *c* and the spike’s main channel), and principal components analysis (PCA) projections **f**_*n*_ of the collision-cleaned waveforms **w**_*n*_. This PCA is fit during initial detection to randomly chosen collision-cleaned waveforms, concatenating all their temporal traces across channels so that the PCA operates only over the temporal axis. Such a “temporal PCA” embedding is stored for each channel of each collision-cleaned waveform. All these features can be used for different steps of the pipeline such as clustering or visualization of detected waveforms along the probe. The detected channel *c*_*n*_ and the detection time *t*_*n*_ are also stored for each event.

### 4.2 Registration

Next, the spike times and localization features are used to estimate probe motion during the recording (*Windolf et al., 2022*). This motion can be the same across the whole probe (rigid) or vary along the depth of the probe (non-rigid). The displacement estimate of the *t*th sample and the *d*th spatial bin is denoted as **P**, a *T* × *D* matrix. In the rigid case, **P** does not change across depth. From here, the registered spike location

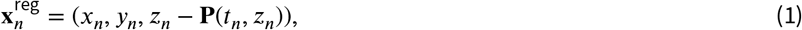

is computed, where **P**(*t, z*) is the displacement estimate at time *t* and depth *z*.

### 4.3 Relocating the waveform features

In order to reduce the dependence of the spike features on the motion of the probe, the feature space should be as drift-invariant as possible. Towards this goal, a model is used to reduce drift in the observation space. In this model, the observed amplitude vector **p***n* for each denoised spike is modeled by the amplitude vector 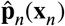 which would arise from a point source at the position **x**_*n*_ (*Boussard et al., 2021*). This point source model minimizes the reconstruction error by finding

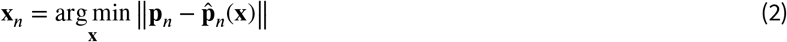

Such a model leads to a natural notion of waveform ‘relocation,’ an operation which changes the source location of a waveform by transforming the observed amplitudes and by shifting the observed channels. Each waveform can be ‘relocated’ to produce a new waveform which appears to originate from the same source at another position. To the extent that the point source model holds, using it to relocate each waveform from its original position to its registered position before computing waveform features such as PCA embeddings will reduce the sensitivity of those features to drift.

To relocate a waveform to correct for drift along the probe’s long axis, the drift is decomposed into two components. The first drift component is determined by the pitch of the probe which is defined as the distance at which the contact geometry repeats along the probe’s long axis (for instance, 40µm for Neuropixels 1.0 probes). This drift component is defined as the maximum multiple of the pitch which is less than the drift. This drift component can be corrected for by shifting the set of channels on which the waveform lies by the corresponding number of pitches. The remaining component of the drift, which is sub-pitch, is corrected for by dividing the waveform at each channel by its predicted amplitude under the point source model at the spike’s location after the first shift and then multiplying by the predicted amplitude at the motion-corrected position.

## 5 Clustering

After the initial detection step, the detected spikes are clustered into putative units. As with many previous spike sorting approaches (*Swindale and Spacek, 2014; Pachitariu et al., 2016; Lee et al., 2020*), the clustering step consists of a rough initial clustering followed by an automated split-merge procedure. The split-merge procedure can then be reused to refine the cluster assignments after receiving a new set of detected spikes from the template matching step (Section 7).

In the following subsections, two strategies are described for initial clustering and split-merge. The first strategy, max-channel assignment, can be utilized for recording devices with relatively sparse densities (e.g. Neuropixels 1.0). The second strategy, global clustering, is more appropriate for extremely high density devices, such as Neuropixels Ultra probes.

### 5.1 Initializing the cluster assignment

#### 5.1.1 Max-channel assignment

For max-channel assignment, the cluster labels *ℓ*_*n*_ are initialized very coarsely to the index of each spike’s registered “max-channel,” defined as the channel closest to its registered position (Eq. (1)). This initial assignment is not expected to be accurate, but instead serves as a way of temporarily dividing the original large dataset into a collection of subsets which can be further divided and refined by the split-merge step below.

#### 5.1.2 Global clustering

For high density probes where the number of channels is equal to or larger than the number of units, max-channel assignment will lead to many oversplits which are hard to resolve with splitting and merging. In such cases, a more effective strategy is to initialize the spike assignments by performing a global clustering of the low-dimensional spike features across the probe. An algorithm that is effective for this task is HDBSCAN∗ (*McInnes et al., 2017*) with a few standard modifications (*Malzer and Baum, 2020*). HDBSCAN is fairly robust with respect to regions of varying spiking density; in addition, it is useful for “triaging away” outlier spikes, similar in spirit to the triaging approach described in (*Lee et al., 2020*). The parameters for HDBSCAN are fixed to `min_cluster_size=min_samples=`15 and `cluster_selection_epsilon =`20μm (*McInnes et al., 2017; Malzer and Baum, 2020*) for all experiments in the paper.

For the global clustering, three features are used per spike: the spike’s horizontal position 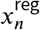, registered depth position 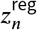 and ptp amplitude *p*_*n*_. The orthogonal location coordinate *y*_*n*_ is not used since, in practice, the estimate of spike’s location along this orthogonal axis tends to be noisy, not providing additional useful clustering information. The amplitude feature is log transformed and rescaled so that its variability is similar to that of the two location features, leading to clusters which are roughly isotropic and, therefore, easier for distance-based clustering algorithms to isolate.

#### 5.1.3 Ensembling initial clustering over time windows

To further improve the global clustering, it is possible to combine clustering outputs from consecutive segments of the recording, aggregating the results with the procedure described in Algorithm 2. As each time window contains observations from a similar distribution of units, this approach can be seen as an “ensembling” method, in which an “ensemble” of multiple clusterings are fused together for improved performance (*Nguyen and Caruana, 2007; Vega-Pons and Ruiz-Shulcloper, 2011*). This approach generally improves the initial cluster assignments (Figure 3), allowing for splitting clusters that are overmerged in one or more of the original time windows. In the presence of drift, this method also allows for tracking units before the template matching procedure. By assigning spikes to each unit at various displacement values, this method provides a first estimate of how each unit’s waveform shape varies with drift. This is leveraged when constructing the superresolved spike train (Section 7). Moreover, if a unit appears or disappears during the recording due to motion or has a low firing rate in most of the windows, it is possible to use the unit’s spikes contained in high firing rate windows to generate a template that can be applied in the low-firing windows.

**Figure 3.**
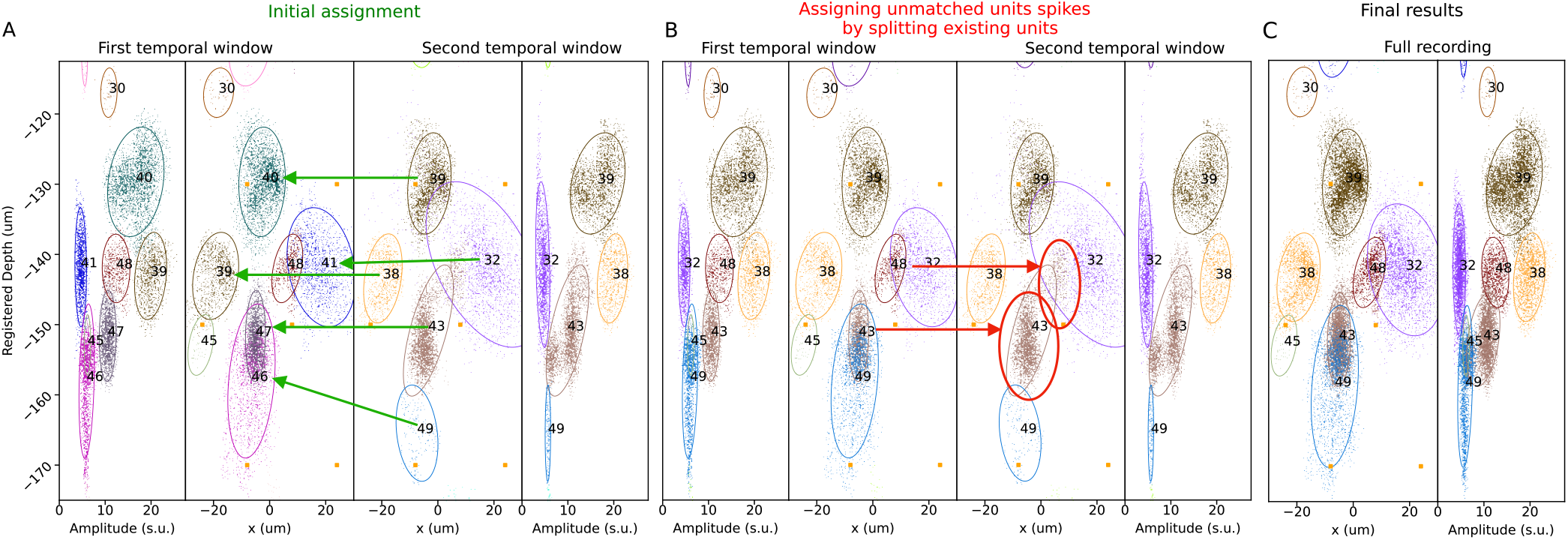
Ensembling HDBSCAN clusterings over time windows in a 600-second MEARec recording (see Section 8 for details). **(A)** The recording is divided into two windows of 300 seconds that are clustered separately using HDBSCAN (see section 5.1.2). In this example, a subsection of the probe is shown for the two temporal windows. From left to right, the four subplots in (A) consist of the x positions and amplitudes of the clustered spikes for the first window, the x positions and registered depths for the first window, the x positions and registered depths for the second window, and the x positions and amplitudes for the second window. This 4-panel view shows the three features used for tracking units over time. To ensemble the clustering over windows, an initial assignment forward pass is done where units of the second window are matched with the closest unit of the first window based on the minimum unit distance (Section 5.1.3). Green arrows represent the matches between units. If several units from the second window are matched with a single unit in the first window, the unit gets split between all its corresponding units. **(B)** Cluster assignments are updated and the labels are matched between both windows. As some units from the first window are not matched by the forward pass (for example, unit 48), a backward pass (red arrows) is run to find the units in the second window closest to the unmatched units. The units in the second window that are matched in this step (for example, unit 43) are split between the first window units. **(C)** After the forward-backward pass, units now contain spikes over multiple time windows.

##### Algorithm 2

Ensembling clustering by tracking units across windows

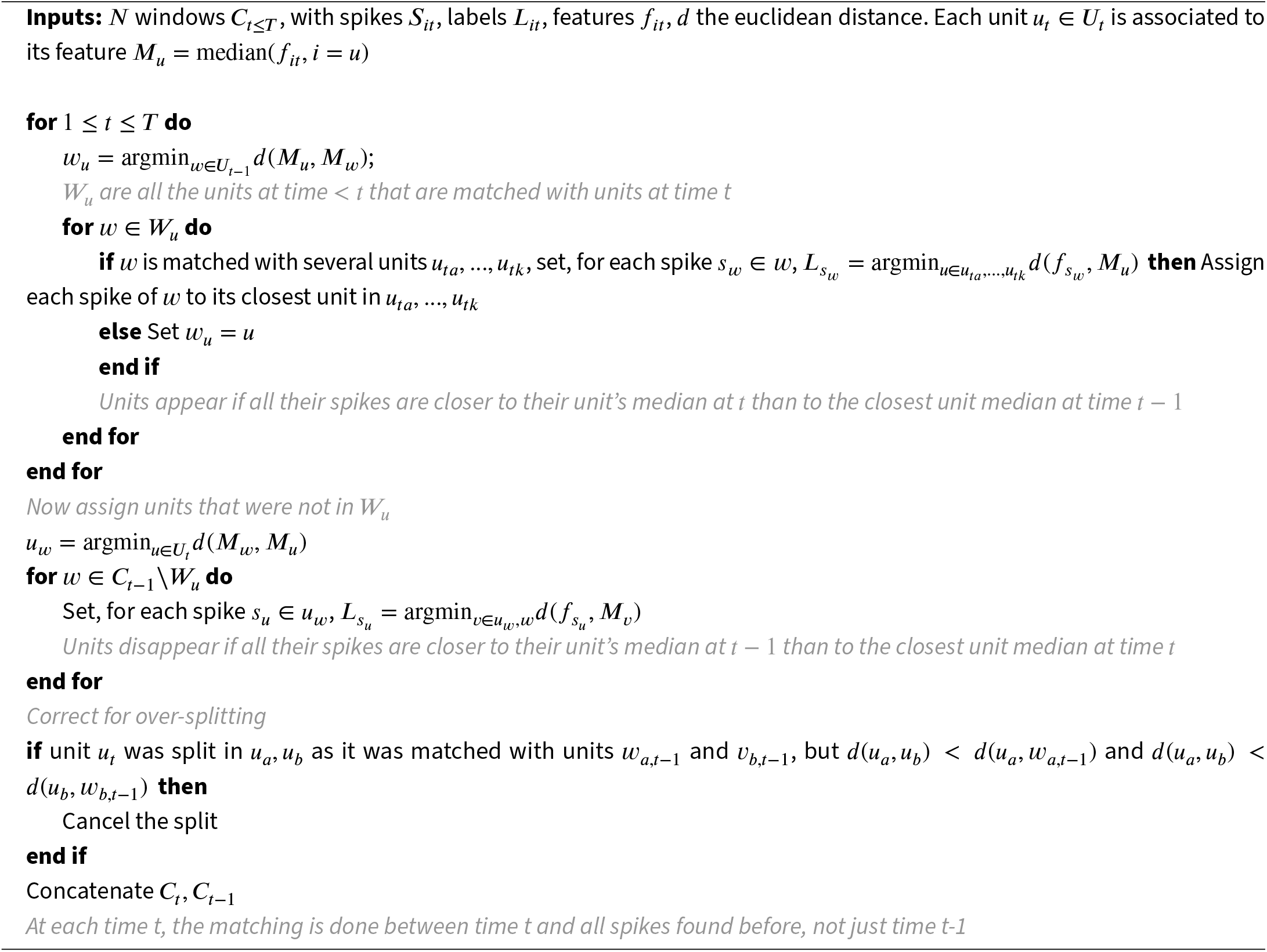

### 5.2 Splitting

The clusters computed in the previous section are found using a highly reduced feature set where the waveform shape is not utilized. This can lead to an initial clustering with too many overmerges or oversplits. To correct for these clustering mistakes, splitting and merging must be performed. Given the set of cluster labels *ℓ*_*n*_, localizations **x**_*n*_, and temporal PCA embeddings **f**_*n*_ of collision-cleaned waveforms **w**_*n*_, the clustering is refined using an automated splitting and merging procedure similar to those of other spike sorters (*Lee et al., 2020; Hennig et al., 2019; Yger et al., 2018; Magland et al., 2020; Chung et al., 2017; Swindale and Spacek, 2014*). The splitting procedure is detailed in this section while the merge step is discussed in section 5.3. The splitting procedure consists of two split steps: (1) recursive HDBSCAN and (2) a diptest-based split.

#### 5.2.1 Splitting by recursive HDBSCAN

This split step is applied to each cluster individually using per-cluster features. When run on a cluster, this step has two potential outcomes: (1) spikes in the cluster are assigned to newly discovered units or marked as outliers, (2) the cluster is kept intact without removing outlier spikes. This step is applied recursively on its own output until no new clusters are discovered.

To apply this split step to a cluster, a set of per-cluster features are computed and then passed into HDBSCAN. HDBSCAN is run using the same parameters as the initial clustering described above. In contrast to the reduced feature set of the initial clustering, the per-cluster set of features now includes additional waveform information. This feature set consists of localization features 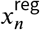 and 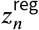 along with features computed as described in section 4.1 using relocated waveforms **w**′ (see section 4.3). The main difference here is that the principal components are computed per cluster and the ptps are extracted from the relocated waveform’s maximum ptp value. The amplitudes are log-transformed and rescaled per cluster such that their standard deviation is equal to that of *x*_*n*_ within each cluster. On highly dense probes, where waveforms are typically seen on hundreds of channels, the first two spatiotemporal principal components may not be informative. In this regime, PCA is fit on the max channel waveform only. To incorporate spatial information into the clustering in this setting, the spread feature *s*_*n*_ is utilized. The spread feature is also rescaled to match the standard deviation of the other features.

#### 5.2.2 Diptest split

In this second split step, the first principal component of the three-dimensional features 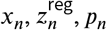 computed per unit. The bimodality of this principal component’s distribution is then tested using the modified Hartigan’s diptest of ISO-SPLIT (*Magland and Barnett, 2016*). This test outputs outputs a statistic close to 0 when the distribution is unimodal and higher than 1 when the distribution is multi-modal. The statistic increases as the probability of unimodality decreases. In the case where there is a multimodal distribution, a cut point can be found which separates the points into each of the two distributions. The spikes are split into two units based on the cut point when the test statistic is higher than a given threshold. This step has the advantage of not labelling spikes as outliers and is used after the initial HDBSCAN clustering and after the final deconvolution. This step was found to be useful for biasing the algorithm towards oversplits in early iterations which is desirable as it allows for detecting as many cells as possible during template matching. These oversplits can then resolved in later processing steps. Therefore, a threshold of 1 is used after the initial clustering step and a higher threshold of 5 is used after the final deconvolution step, where only “obvious” overmerges should be corrected.

### 5.3 Merge step

#### 5.3.1 Obtaining candidate merge pairs

As testing for merges across all pairs of units is too computationally expensive, the merge step only tests a subset of possible unit pairs. Three methods are proposed for selecting candidate pairs. The first method relies on a template distance. The templates are first computed for each unit then deconvolution (Section 7.3) is run from each unit’s template on all other units’ templates. Candidate pairs are proposed when the deconvolution residual’s maximum absolute value is lower than a threshold as the two templates are similar and were likely generated due to over-splitting. Second, templates can again be computed for each unit and candidate pairs proposed when templates for two units both share large amplitude on at least one common channel. Third, the units’ distances in the registered space can be utilized to propose candidate pairs. If the median registered position of a unit’s spikes is less than one pitch of the probe from another unit’s median registered position, a merge is proposed for these units.

#### 5.3.2 Drift-aware template merge

A drift-aware merge step is applied to all candidate merge pairs. Superresolved templates (Section 7.1) for each unit are first computed. As described in Section 7, each unit will have multiple templates which correspond to different depth “bins” in order to capture variability due to drift. The superresolved templates are then shifted by the difference between the depths of locations where the templates have been computed and, for “overlapping bins”, the templates of the first unit are deconvolved with the templates of the second, shifted unit following the procedure described in Section 7. For example, if two units *u*_1_, *u*_2_ are proposed for a merge, and located around median depth *z*_1_, *z*_2_, the super-resolved templates of *u*_1_ are shifted by *z*_2_ − *z*_1_. Then, for bins around *z*_2_ that contain superresolved templates for both units, the templates of *u*_2_ are deconvolved with the shifted templates of *u*_1_. The L2-norm of the residual of deconvolved templates is then computed and divided by the L2-norm of the first unit’s templates. The resulting quantity is dimensionless so a single threshold can be set to compare distances between templates. This distance, and the corresponding optimal offsets, can be computed efficiently for all candidate pairs. Since each pair will have two distance values, the maximum of these two values is chosen as the distance between the units.

Once the above distance is computed for all pairs of units, candidate merge pairs are merged by running an agglomerative clustering on the corresponding distance matrix. A distance threshold of 0.25 is typically used to merge units, but this parameter can be adjusted depending on the goal of the current processing step. For example, it is possible to use a smaller value 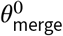to encourage oversplitting and template diversity in earlier split-merge steps before increasing it to a larger threshold 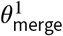at the final merge step. This process is illustrated in Figure 4.

**Figure 4.**
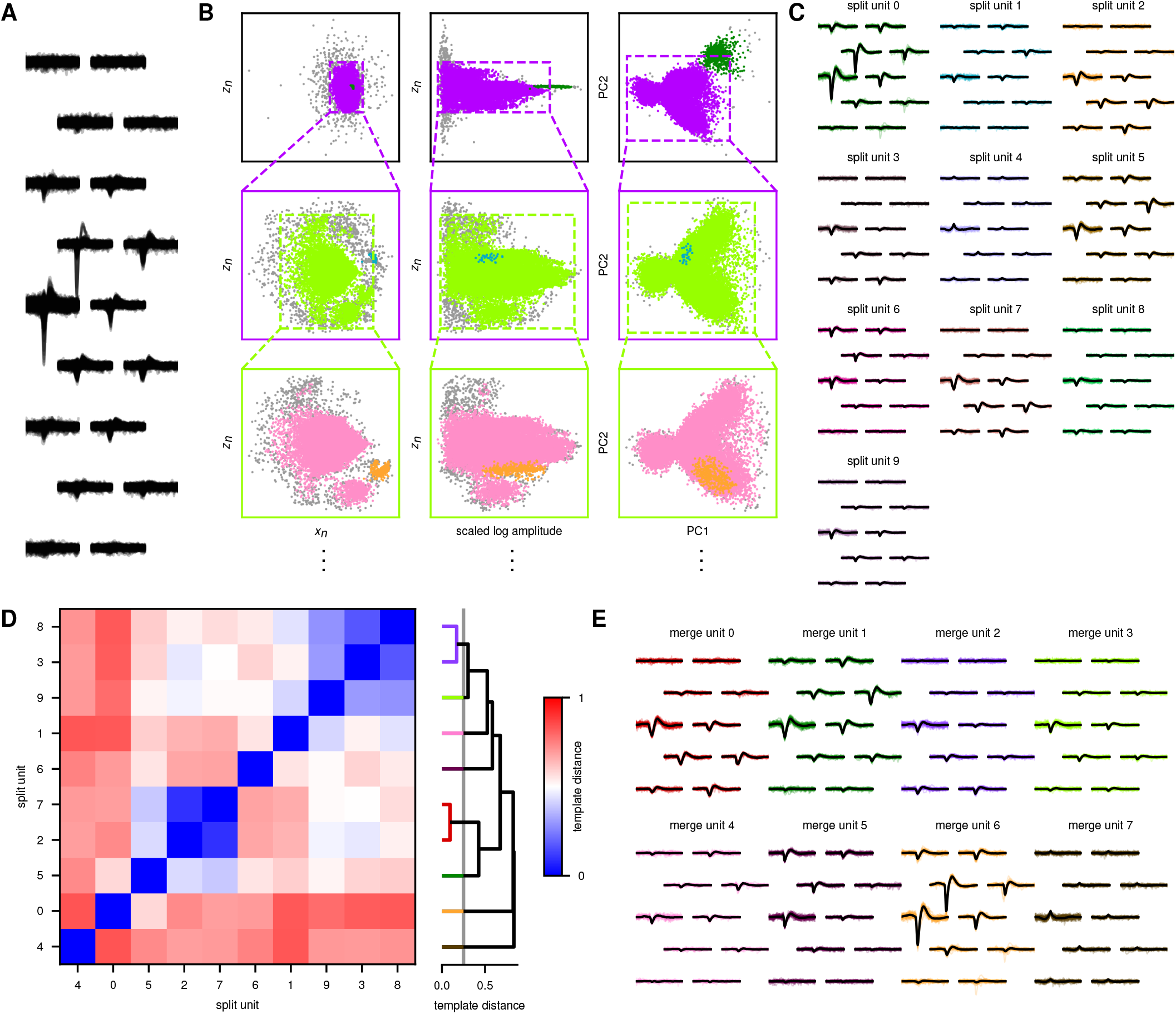
Overview of the recursive clustering step. In this figure, splitting and merging steps are applied to waveforms detected on a single channel in an IBL recording (*IBL et al., 2022*), demonstrating a simplified version of the split-merge with max-channel assignment described in Section 5. **(A)** shows a random collection of cleaned waveforms **w**_*n*_ on a channel neighborhood around the detection channel. In **(B)**, location (left column), amplitude (center), and per-cluster PCA features (right) are recursively clustered with HDBSCAN. First, clustering is done on the full collection of waveforms detected on the channel (top row). Then, HDBSCAN is performed on each newly discovered cluster (middle, bottom) until no new clusters are found. Representative waveforms and denoised templates (Section 6) from the split clusters are shown in **(C)**. In **(D)**, the normalized, dimensionless template distance between units is shown together with the complete-linkage dendrogram used to merge the split units (the merge threshold of 0.25 is shown vertically in gray). This merge step is detailed in section 5.3. Representative waveforms and denoised templates for the resulting merged units are shown in **(E)**. Colors are shared within (but not across) rows (A-C) and (D,E).

This merge step is conceptually similar to other template distance-based approaches such as those used by *Lee et al*. (*2020*) or *Chen et al*. (*2021*). The main advantage of this proposed approach is that it is drift-aware and dimensionless.

## 6 Denoised templates

Given a set of spike times and putative unit labels, the next step is to compute representative template waveforms. For each unit, a template is computed in each superresolved bin (see Section Section 7.1) rather than across all spikes in the unit. Typically, the mean or median of the unit’s raw waveforms is used as the template. Some spike sorters model variability within units (*Garcia and Pouzat, 2015; Yger et al., 2018*) and others consider low rank reconstructions of templates (*Lee et al., 2020; Pachitariu et al., 2016*). In all these cases, such estimators have high variance for units with few spikes, leading to noisy templates.

This issue can be alleviated by exploiting the similarity of waveform shapes for all neurons across the recording. To this end, the template for each unit (or superresolved bin) is modeled as a weighted average of its median raw waveform and the median of its PCA-denoised waveforms, where the PCA is fit to waveforms from all detected units. (Simple PCA applied directly to the collection of templates led to worse results, flattening important differences between the templates, particularly near their peaks.) For the weighting, a hand-designed function is constructed which varies over time and across channels. Intuitively, the weight placed on the raw template should increase as the signal-to-noise ratio (SNR) of the template increases, and the population-based low rank model should be emphasized when this SNR is small. We choose to approximate this SNR for each channel by the template’s amplitude on that channel multiplied by the square root of spike count, since in a standardized recording the variance of the sample mean will be inversely proportional to the spike count. The weighting function combines this notion of per-unit and per-channel SNR with a factor that decays to 0 at the temporal edges of the waveform, so that the raw median is emphasized most near the spatiotemporal peak. Visualizations of this weighting, the raw median and low-rank templates, and the significant improvement in accuracy of this method are displayed in Fig. 5.

**Figure 5.**
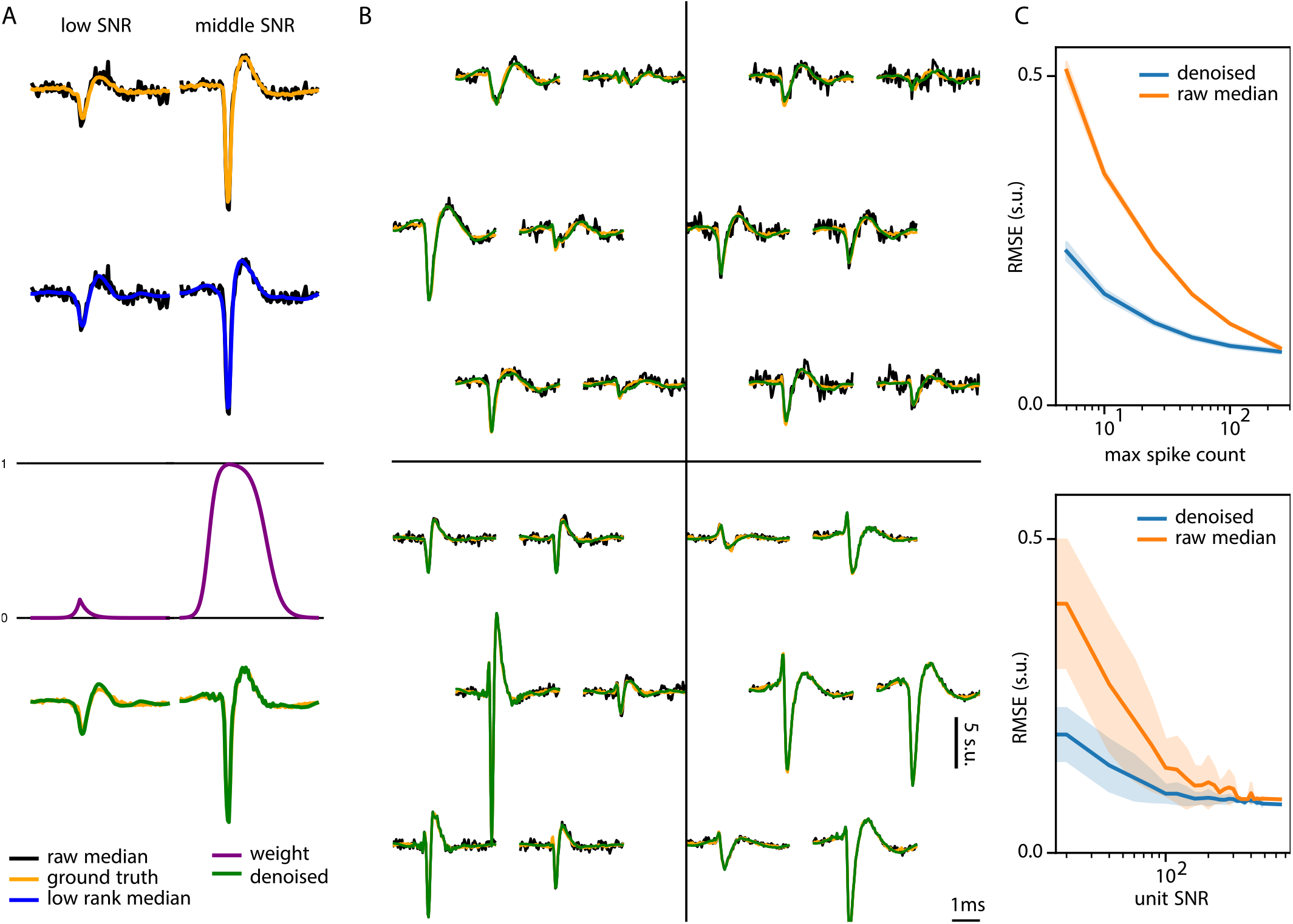
A hybrid data study of the template denoising approach described in Section 6. Templates collected from previous spike sorting runs (*IBL et al. (2022*)) were added into snippets of raw data, allowing for comparisons between the denoised templates and the standard median templates in terms of their distance (RMSE) from the ground truth (GT) template. In **(A)**, from top to bottom: GT templates (orange) overlaid over raw median templates (black); low-rank templates (blue) overlaid over raw median templates; the temporal mixing weight (purple), where a higher weight means that the raw median will makeup a larger portion of the denoised template at this sample; GT templates overlaid over denoised templates (green). In **(B)**, raw, denoised, and GT templates are overlaid for four units across small channel neighborhoods around the templates’ main channel. In **(C)**, the RMSE of the denoised templates is compared to that of the raw median templates. This quantity is plotted as a function of the number of spikes used to compute the template and the SNR (computed as the amplitude multiplied by the square root of spike count). The denoised templates have uniformly lower RMSE than the raw median templates especially for lower SNRs and spike counts. Error bars are the standard deviations of all 100 injected units’ RMSEs across three randomly generated hybrid recordings.

## 7 Drift-tracking template matching algorithm

The previous steps output cluster labels that assign detected spikes to putative units. This allows for constructing representative template waveforms for each unit. The goal of a template matching or deconvolution algorithm is to use these templates to recover additional spiking events that belong to each cluster. This fills the role of the expectation step in an expectation-maximization clustering algorithm. Iterating deconvolution and split/merge steps tend to improve the assignment accuracy (see Figure 8C below).

In the case where a recording contains drift (approximated by the drift estimate P), naively computing a single template per cluster is insufficient. This is because each unit’s waveforms will be spread across many different positions along the probe such that a naive average would be blurred and shrunk compared to the actual waveforms, leading to errors in the template matching step. To address this, several ‘superresolved’ templates per unit can be computed to capture variability due to drift. These can then be shifted to match the drift estimate as part of a template matching procedure.

To this end, as in (*Pillow et al., 2013; Pachitariu et al., 2016; Lee et al., 2020; Yger et al., 2018; Garcia and Pouzat, 2015; Diggelmann et al., 2018*), the recording **V** ∈ ℝ^*T* ×*C*^, which has *T* time samples and *C* channels, is modeled as the sum of a convolved spike train **X** and a residual **R**:

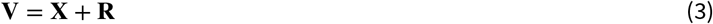

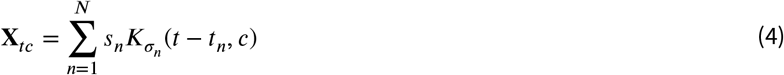

Here, *N* is the total number of spikes, *t*_*n*_ is the spike time of the *n*th spike, and *s*_*n*_ ∼_iid_ *N*(1, λ) are scaling terms allowing for amplitude variability. The main difference of this proposed algorithm is that σ_*n*_ is not the unit label *ℓ*_*n*_ of each spike, but rather an index into a collection of templates which have been grouped together for each putative unit in a drift-aware manner as well as temporally upsampled and shifted according to the drift. The construction of these templates and how they are used in the drift-tracking template matching algorithm is discussed in detail below (see also Figure 6).

**Figure 6.**
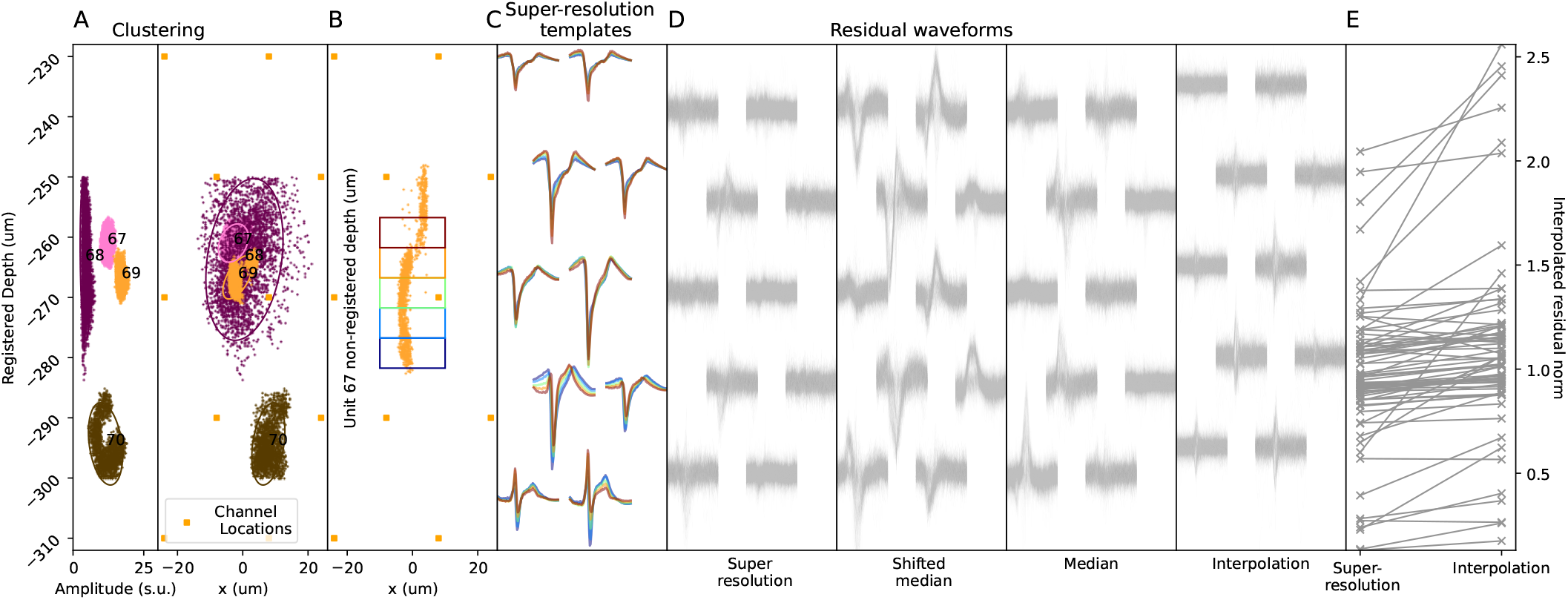
Superresolved templates lead to better matching between templates and spikes from the same unit in simulated data. **(A)** shows the registered depth vs. the amplitude (left panel) and horizontal location x (right panel) of 5 clusters found after initial clustering of a 600s MEARec simulated recording with induced motion (see section 8 for details). **(B)** shows the unregistered positions of cluster 69’s spikes in pink with the depth coordinate divided into 5 bins of size 5um. Five superresolved templates are computed using the spikes in each of these bins. **(C)** shows the superresolved templates for cluster 69 on 10 channels. Each superresolved template is colored by the same color as the bin it corresponds to in panel (B). The superresolved template corresponding to the inferred position of each unit can be used to capture the unit’s spike shape variability at each time step. **(D)** shows the residuals for a unit after subtracting templates generated using various template generation schemes. From left to right, the panels show the residuals obtained using superresolved templates, median shifted templates, median templates, and interpolation (used by Kilosort 2.5). Panel **(E)** shows, for every unit, the norm of the mean squared residuals for the superresolution templates versus those of the interpolation templates. Each connected pair of points corresponds to a single unit. The superresolution templates have a lower average norm of the mean squared residual than the interpolation templates.

### 7.1 Constructing the superresolved templates

To capture the variability within each unit due to drift, each unit is spatially ‘superresolved,’ i.e., the unit is broken into several sub-units based on its movement along the probe. First, a bin size is set such that it is a fraction of the probe’s pitch (typically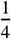or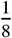 of the pitch). Within each unit, spikes are binned according to their unregistered depths *z*_*n*_ such that each spike is assigned to a superresolved bin. Then, denoised templates are computed for each unit (see section 6) using the spikes in each superresolved bin. If a bin has too few spikes (in practice, fewer than 25), spikes one pitch away in the corresponding bin are utilized instead. This procedure theoretically allows for computing templates at every position along the probe for each unit, but in practice templates are only computed for bins that cover the registered depth spread of each unit. This still allows for modeling each unit’s shape variability at all timepoints.

Figure 6A-B show an example of how superresolved templates are constructed for one unit. Figure 6C shows the different versions of the superresolved templates. Figure 6D shows the residuals for one unit after subtracting templates produced by four different template generation schemes. The first panel shows the residual after subtracting the superresolved templates from the raw waveforms, the second after subtracting the median template shifted according to the displacement, the third after subtracting the median template without any shift, and the fourth after subtracting a median template extracted after interpolation to correct for drift. For the interpolation, a kriging interpolator is used (*Steinmetz et al., 2021*). The residuals left by the superresolved templates contain less signal that those of the interpolated data templates.

Even though residuals might appear smaller in the last panel, interpolating the raw data to correct for drift also reduces SNR, as both residuals and spikes will be smaller. In Figure 6E, the norm of the superresolved template residuals versus the norms of the interpolated template residuals are shown for each unit. For a fair comparison, the residuals of the superresolved templates are interpolated before computing the norm, which is equivalent to interpolating the superresolved templates before subtracting the raw data (interpolation is a linear operation). A significant decrease in the residual norm (average decrease of 19%, p-value 0.04) is obtained when using superresolved templates rather than interpolated data templates, illustrating how superresolved templates capture motion-induced variability while avoiding any SNR loss or interpolation artifacts.

### 7.2 Shifting the superresolved templates to track large drift

Given a set of superresolved templates computed for each unit, it is straightforward to extend basic template matching algorithms to handle drift. Since the set of superresolved templates is computed according to the drift, these templates will contain the shape variability of each unit as it drifts across the probe. In other words, the superresolved templates model the sub-pitch drift.

To model larger scale drift, the position of each unit over time is estimated as *z*_*u,t*_. As each unit’s superresolved templates are computed around a depth *z*_*u*,comp_, they need to be shifted by *z*_*u,t*_ −*z*_*u*,comp_ to model that unit’s shape at time *t*. For each unit *u* with bin size *h*_*z*_ and pitch Δ_*z*_, the number of pitch shifts *k* = (*z*_*u,t*_ − *z*_*u*,comp_) mod (Δ_*z*_) and the sub-pitch shift *b* = (*z*_*u,t*_ − *z*_*u*,comp_ − *k* ∗ Δ_*z*_)(mod *h*_*z*_) are computed. The superresolved templates are then shifted by *k* pitches before shifting *b* of these superresolved templates by one pitch. Note that this shift is different from the relocation described in Sec. 4.3, as we take advantage of multiple waveform localization in each cluster here.

These superresolved templates are then passed as input into the template matching subroutine which yields a new spike train. Spikes that are assigned to any of the superresolved templates from a unit are then assigned to that unit.

### 7.3 Template matching procedure

Similar to prior work (e.g., (*Pachitariu et al., 2016; Lee et al., 2020*), given a set of *k* templates *K*_1_, …, *K*_*k*_, a greedy template matching algorithm is used to optimize the following cost (with 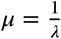):

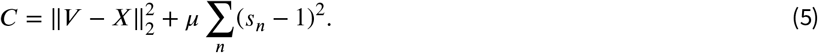

As defined in Eq. (4), **X** is the convolved spike process or, in other words, the sum of the scaled and temporally subsampled templates *s*_*n*_*K*_σ*n*_ . Here, σ_*n*_ is an index into a collection of temporally upsampled versions of the *k* original templates. The algorithm proceeds with a coarse-to-fine approach, selecting the best matching template of the *k* original templates at the best temporal position in the recording and then refining this match by finding the best temporally upsampled version σ_*n*_ of the template and the optimal scaling value *s*_*n*_.

Coarse matches are found by optimizing the decrease in the cost function due to adding a spike of amplitude 1 from unit *u* at a temporal offset *t*. This decrease is provided through a *K* × *T* matrix,

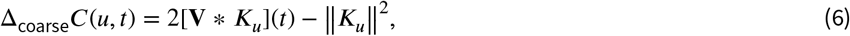

where the cross-correlation ∗ is a temporal filtering followed by a sum over channels. The cross-correlation operation is accelerated using a low-rank approximation of the templates, following (*Pachitariu et al., 2016*).

Then, after a coarse match is found, the scaling and temporal upsampling are determined by optimizing the fine cost function, i.e., the cost of adding a spike from the upsampled templates σ at time *t* for the optimal scaling *s*

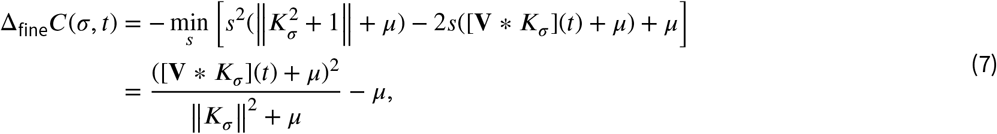

where the optimal scaling is achieved by

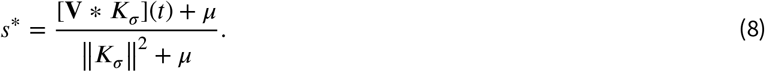

Again, following (*Pachitariu et al., 2016*) and other previous matching-pursuit approaches, the cost functions are updated directly (and in parallel) rather than updating the original data residual. This procedure is iterated until there is no spike which decreases the cost by more than some threshold value Δ_coarse_*C*(*u, t*) ≥ θ_deconv_.

## 8 Evaluation on synthetic datasets

In this section, we benchmark DARTsort and Kilosort 2.5 (with default parameters) on synthetic datasets with multiple drift types and varying noise levels. We show that DARTsort outperforms Kilosort 2.5 on all drifting datasets and on non-drifting datasets with increased noise levels, demonstrating that DARTsort is more robust to drift and noise.

### 8.1 Synthetic datasets

Nine 600 second long simulated datasets were generated using the extracellular simulation software, MEArec (*Buccino and Einevoll, 2021*). Each synthetic dataset is based on a subset of Neuropixels 1.0 geometry, consisting of 64 electrodes recording at 30kHz. Spikes are simulated from 64 units which are distributed around the array uniformly and randomly.

For these datasets, three different drift types are included, (1) static (i.e., no drift), (2) rigid zig-zag motion, and (3) rigid ‘bumps’ (Figure 7A). The zig-zag motion dataset is comprised of slow up-and-down drift across the whole probe starting after a fixed time period. The ‘bumps’ data consists of periods with no motion interspersed with sudden motion across the whole probe. For each drift type, three datasets are simulated with increasing noise levels. MEArec’s distance-correlated noise mode is used, which consists of a multivariate Gaussian with a covariance matrix that decays as a distance from the main channel.

**Figure 7.**
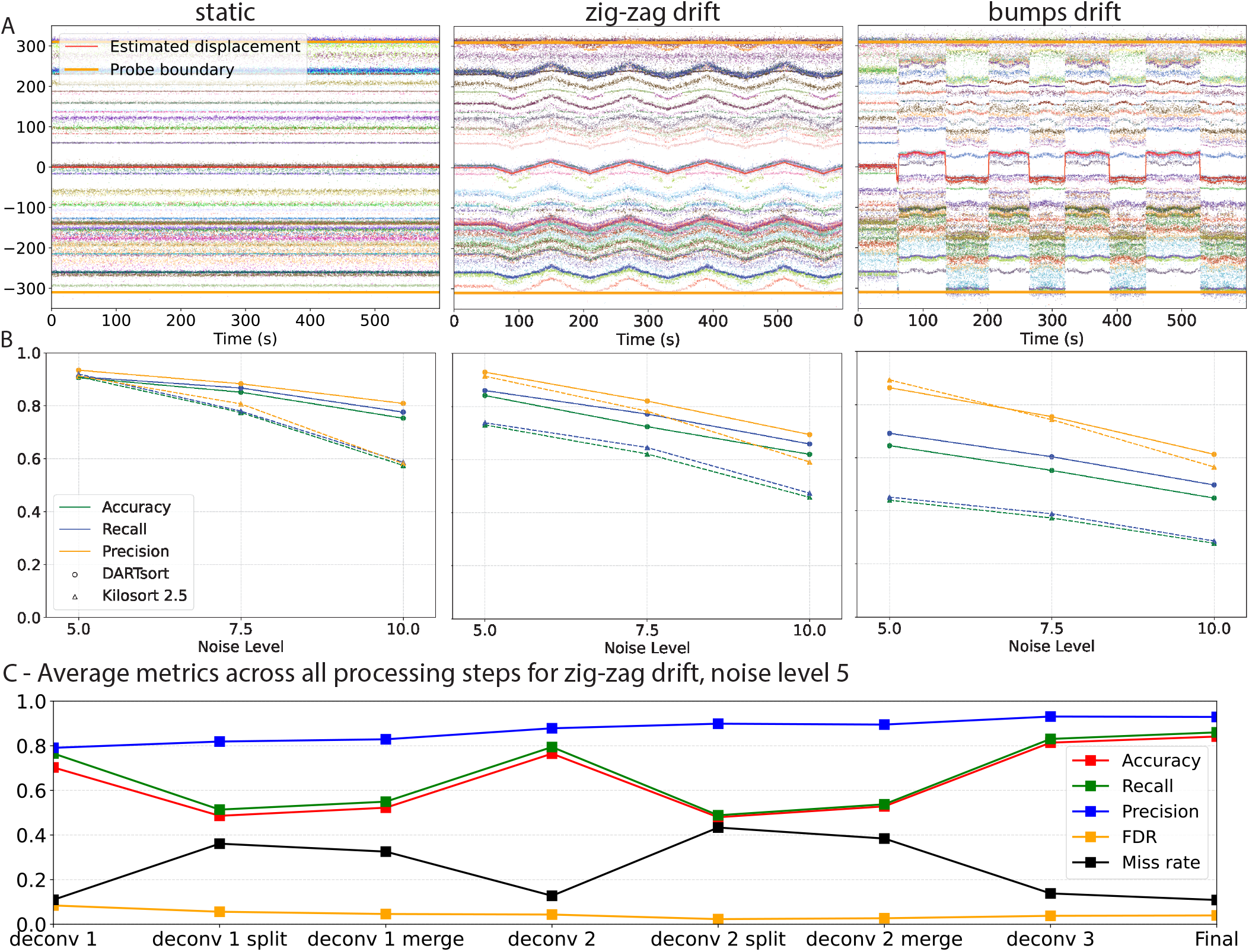
DARTsort outperforms Kilosort 2.5 on synthetic data. **(A)** The raster maps for DARTsort on the lowest noise MEArec dataset (noise level 5 - see section 8). The estimated displacement of DARTsort is overlaid at depth zero (red trace) and the probe boundaries are overlaid at the top and bottom of the figure (orange lines). **(B)** The accuracy, recall, and precision for all datasets for DARTsort and Kilosort 2.5. DARTsort outperforms Kilosort 2.5 across noise levels. **(C)** The accuracy, recall, precision, false-discovery rate (FDR), and miss-rate for an example MEArec dataset across all steps of DARTsort’s processing pipeline. The precision consistently increases across all stages of the pipeline while the recall is improved during each merge step and deconvolution. The miss rate increases during each split step due to outlier triaging by the HDBSCAN split step. By the end of the pipeline, the accuracy is maximized and the FDR and miss rate are low.

**Figure 8.**
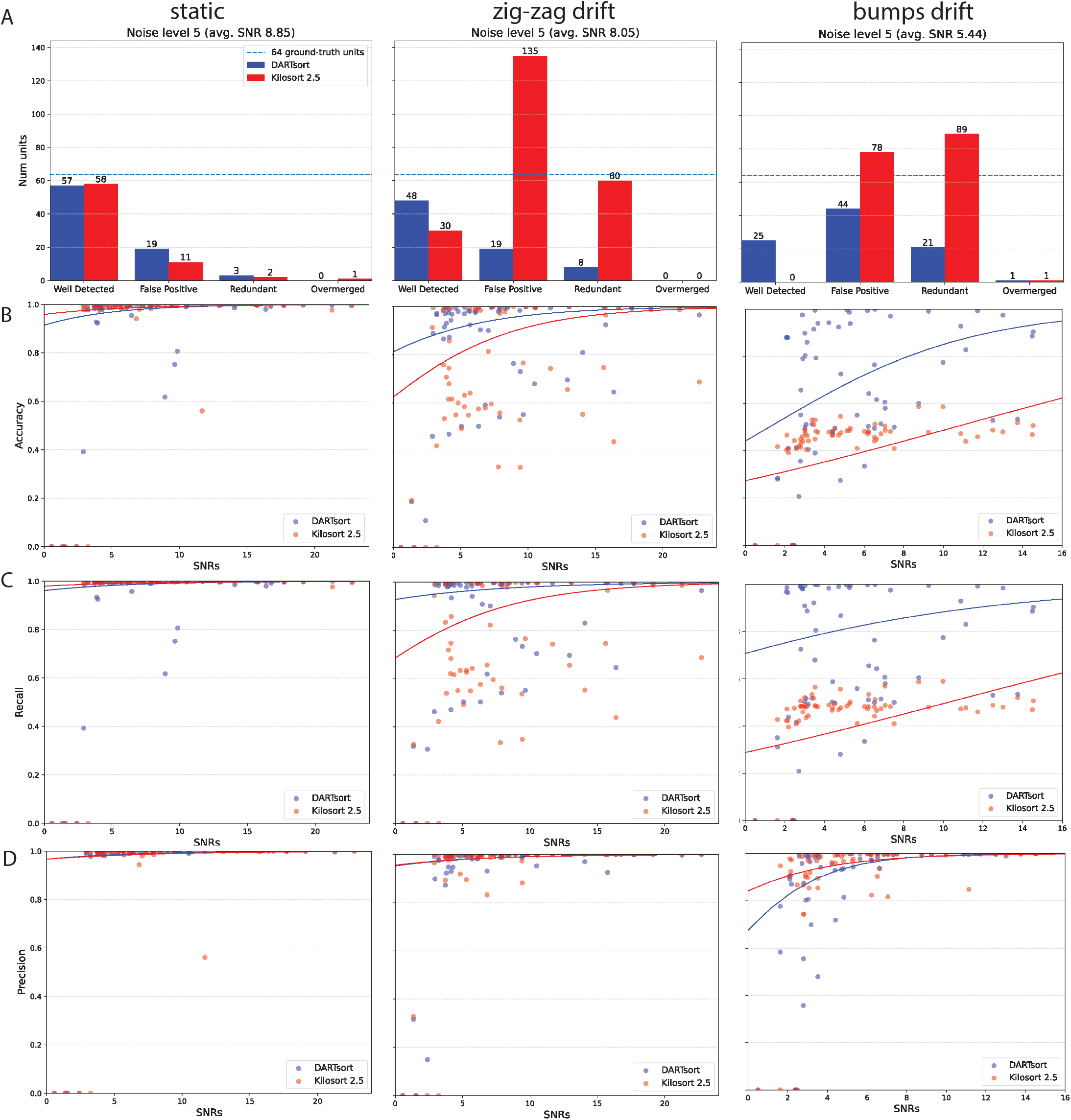
DARTsort comparison to Kilosort 2.5 on low noise simulated data. **(A)** The unit summary of DARTsort and Kilosort 2.5 for noise level 5. Results are similar for the static dataset. DARTsort has more well-detected units and fewer bad units (false-positive and redundant) than Kilosort 2.5 on both drift types. **(B)** The signal-to-noise vs accuracy of each of the 64 ground-truth units from the noise level 5 dataset. Each dot indicates a single unit; trace indicates logistic fit. DARTsort recovers more low SNR units than Kilosort 2.5 on both drifting datasets while still performing better on high SNR units. **(C)** and **(D)** show the precision and recall of DARTsort and Kilosort 2.5, respectively, for the three datasets. For all datasets, DARTsort’s recall is similar to or better than Kilosort 2.5. For the precision, the two methods are similar.

DARTsort was run using default parameters as described in the text. The neural-network denoiser (Section 4) was trained previously on templates extracted from NP1 recordings (*IBL et al., 2022*) and not on MEArec templates. Also, while template denoising (Section 6) is useful when processing real extracellular datasets, it is not used for these simulated results.

### 8.2 Results

To evaluate the performance of DARTsort and Kilosort 2.5, SpikeInterface’s ground-truth comparison module is utilized. Both sorters are evaluated on their precision, recall, and accuracy on the 64 ground-truth units. The sorters are also evaluated for the number of additional units they find, as false positive units are a challenge for interpretation and postprocessing. The unit categorization scheme of well-detected, false positive, redundant, and overmerged units are used in these results. For more details about these categorizations, see (*Buccino et al., 2020*).

The results for all synthetic datasets are are shown in Figure 7. The accuracy, recall, and precision for all datasets are shown in 7B. For the static datasets, although the performance of DARTsort is similar to Kilosort 2.5 in the lowest noise regime, DARTsort is better when the noise is increased. At the highest noise level, DARTsort’s performance is still high (75% average accuracy) while Kilosort 2.5’s performance dips significantly (57 % average accuracy). For the drifting datasets, DARTsort outperforms Kilosort 2.5 across all noise levels. These results demonstrate that DARTsort is more robust to drift and noise than Kilosort 2.5 on simulated data.

Figure 7C shows the accuracy, recall, precision, false-discovery rate (FDR), and miss rate for an example synthetic dataset across all steps of DARTsort’s processing pipeline. The precision of DARTsort consistently increases across all stages of the pipeline, while the recall is improved with each merge and deconvolution step. The miss rate increases and the recall decreases during each split step as spikes are triaged away by the recursive HDBSCAN split step.

Per-unit metrics are shown across three low noise, synthetic datasets with different drift types in Figure 8. Figure 8A shows unit summary bar plots for each unit outputted by DARTsort and Kilosort 2.5. DARTsort significantly outperforms Kilosort 2.5 on both of the drifting datasets, finding 48 vs 30 and 25 vs. 0 well-detected units on the zig-zag and bumps dataset, respectively. Crucially, DARTsort also outputs significantly fewer false-positive (19 vs. 135 and 44 vs. 78) and redundant units (8 vs. 60 and 21 vs. 89), facilitating post hoc interpretation and curation.

In Figure 8B-D, the SNR of each unit is plotted vs. the accuracy, recall, and precision of each sorter, respectively. The SNR is computed as the maximum amplitude of average spike waveform on the best channel divided by the median absolute deviation of the background noise on the same channel. For all datasets, DARTsort has similar or better recall than Kilosort 2.5 and comparable precision except for on the bumps dataset (where DARTsort’s recall is much higher). Also, DARTsort is able to recover more low SNR units than Kilosort 2.5 while maintaining a low false-positive rate.

## 9 Future work

A straightforward extension to DARTsort is allowing for non-rigid displacement in the deconvolution. Although the superresolved deconvolution described in 7.3 will work in this non-rigid regime, the current approach relies on a precomputed set of cross correlations between the shifted, superresolved templates. As the number of shifted, superresolved templates will be much higher due to the non-rigid drift, a strategy based on updating the residual (rather than updating the pre-computed cross-correlations) will likely be more efficient.

Along with improvements to the algorithm, more evaluation must be performed to determine if DARTsort is more robust to drift and noise than other spike sorters on both real and hybrid datasets.

## 10 Acknowledgements

We thank Alessio Buccino, Gaelle Chapuis, Jennifer Colonell, Margot Elmaleh, Samuel Garcia, Kenneth Harris, Ari Pakman, Nick Steinmetz, Erdem Varol, Pierre Yger, and also the Simons Collaboration on the Global Brain Spike Sorting Working group for many useful discussions.

